# Whole-genome sequencing of CRFK and PG-4 cells to infer the phenotype of the original donor cats

**DOI:** 10.1101/2025.10.31.685720

**Authors:** Ganma Tanaka, Rikuto Goto, Tetsushi Komoto, Akiko Kubota, Reo Hayashi, Takeshi Igawa, Naoaki Sakamoto, Akinori Awazu

**Author notes:** Ganma Tanaka and Rikuto Goto contributed equally to this work and share first authorship. **Corresponding author:** Akinori Awazu, Graduate School of Integrated Sciences for Life, Hiroshima University, 1-3-1, Kagamiyama, Higashi-Hiroshima 739-8526, JAPAN.

## Abstract

**Background:** Crandell-Rees Feline Kidney (CRFK; kidney-derived) cells and PG-4 cells (astrocyte-derived) have been in use and have been passaged for decades in laboratories worldwide; however, no detailed information on the genetic background of the donor individuals is available, particularly regarding phenotype characteristics such as coat and iris color.

**Results:** We performed whole-genome sequencing of CRFK and PG-4 cells. We analyzed the resulting data to infer the phenotype of the individual from which the cells were derived, specifically for the coat color, coat length, coat pattern, and iris color. Our data suggested that CRFK cells originated from a cat with long black fur lacking stripes and with non-blue irises; PG-4 cells originated from a cat with long bicolored white and black fur without stripes, and with non-blue irises. Analysis of publicly available RNA-seq data confirmed that genes associated with coat phenotype and iris color are expressed in the skin and eyes, as well as in various other organs.

**Conclusions:** Variants of the genes affecting coat phenotype and iris color may influence physiological functions throughout the body. These results shed light on the previously unknown genetic background of commonly used feline cultured cells and the phenotype of the donor individuals. These findings may facilitate more accurate interpretation of data obtained from cultured feline cells and provide guidelines for developing cell lines with domestic cat genotypes exhibiting diverse phenotypes through genome editing. This will help to provide clarity regarding deafness in cats with white fur and blue irises and elucidate the effects of melanocyte destruction caused by KIT gene mutations on the nervous system and the influence of other coat-related factors in organs other than the skin or functions independent of coat formation.

## Background

Domestic cats (*Felis catus*) are widely kept as companion animals around the world, and the phenotypic diversity of their coat color, coat pattern, coat length, and iris color has been a topic of interest to breeders and scientists alike. Recent advances in genomics have markedly improved our understanding of the genetic basis controlling these diverse phenotypes, and the respective genotype-phenotype relationships are being increasingly elucidated.^1^ Examples include the *TYR* gene, which is the color locus associated with albinism and a color point phenotype^2–4^; the *TYRP1* gene, conveying brown coat color^2,5^; the *MC1R* genes, associated with in red coat color^6^; the *KIT* gene, determining white coat and white spotting^7–11^; the *ARHGAP36* gene, involved in orange coat color^12,13^; the *MLPH* gene, causing dilute pigmentation^14^; the *ASIP* gene, controlling the expression of the agouti pattern^15^; the *LVRN* gene, determining the type of tabby pattern^16^; the *DKK4* gene, associated with the ticked tabby pattern^17^; the *FGF5* gene, involved in long-fur variants^18,19^; the *KRT71* and *LPAR6* genes, involved in hairless and rexing (curly hair) phenotypes^20–22^; and the *PAX3* gene, which is associated with blue irises^23,24^. Furthermore, technological advancements such as long-read sequencing facilitated the construction of a more continuous and accurate draft genome of the domestic cat ^25–27^, and a plethora of genetic polymorphisms associated with disease have been identified ^17,25,28–32^.

Immortalized cell lines are an indispensable tool in current biomedical and veterinary science. Crandell-Rees Feline Kidney (CRFK) cells, derived from the kidneys of a domestic cat^33–34^, have been used for numerous virological studies, including feline calicivirus,^35–37^ feline immunodeficiency virus^38–41^, feline coronavirus^42–45^, and various feline endogenous retroviruses such as RD114^46–48^, as well as for vaccine development^49–51^. PG-4 cells, derived from feline fetal astrocytes, have also played an important role in retrovirus research^52–53^.

Additionally, Crandell et al.^33^ reported that CRFK cells exhibited aneuploidy (2n = 37) compared with the normal feline autosome count of 2n = 38, and demonstrated a characteristic karyotype with three marker chromosomes. Lasfargues et al.^54^ also observed that the chromosome count fluctuated between 36 and 38 in each cell. By contrast, no such findings have been reported for PG-4 cells.

These cell lines have been passaged for decades in laboratories worldwide and have facilitated numerous scientific breakthroughs. However, despite their widespread use, no detailed information on the genetic background of the donor individuals, particularly with regard to their phenotype, such as coat and iris color, is available. Genomic information on such cell lines is crucial for a deeper understanding of their biological properties, and elucidating the phenotypes of their donors may provide new insights into the influence of specific genotypes on cell properties.

In this study, we performed whole-genome sequencing of CRFK and PG-4 cell lines. The sequencing data were used for in-depth analysis of the nucleotide sequences of genes known to be associated with coat and iris phenotypes (i.e., *TYR*, *TYRP1*, *MC1R*, *KIT*, *ARHGAP36*, *MLPH*, *ASIP*, *LVRN*, *DKK4*, *FGF5*, *KRT71*, *LPAR6*, and *PAX3*; Table 1). Based on the genotyping results for each gene, we inferred the previously unknown phenotypes of the donor individuals. Furthermore, analysis of publicly available RNA-seq data confirmed that these genes were expressed in skin and eyes as well as in various other tissues throughout the body. The findings and genome sequence data are expected to provide fundamental insights into the genetic background of extensively used feline cell lines. This information will support the interpretation of future research outcomes involving these cell lines and contribute to the development of guidelines for establishing cell lines with various genotypes through genome editing.

**Table 1:**
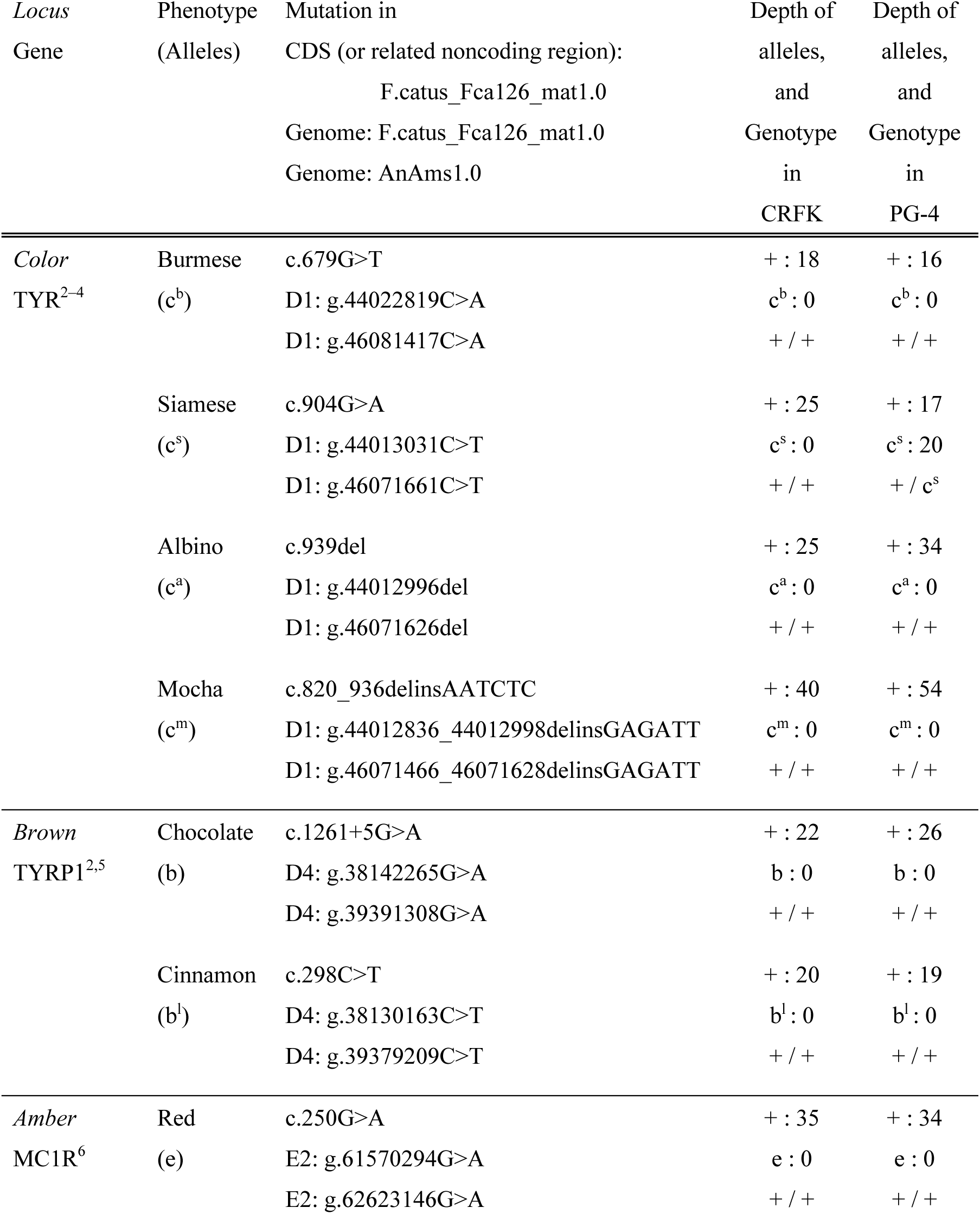
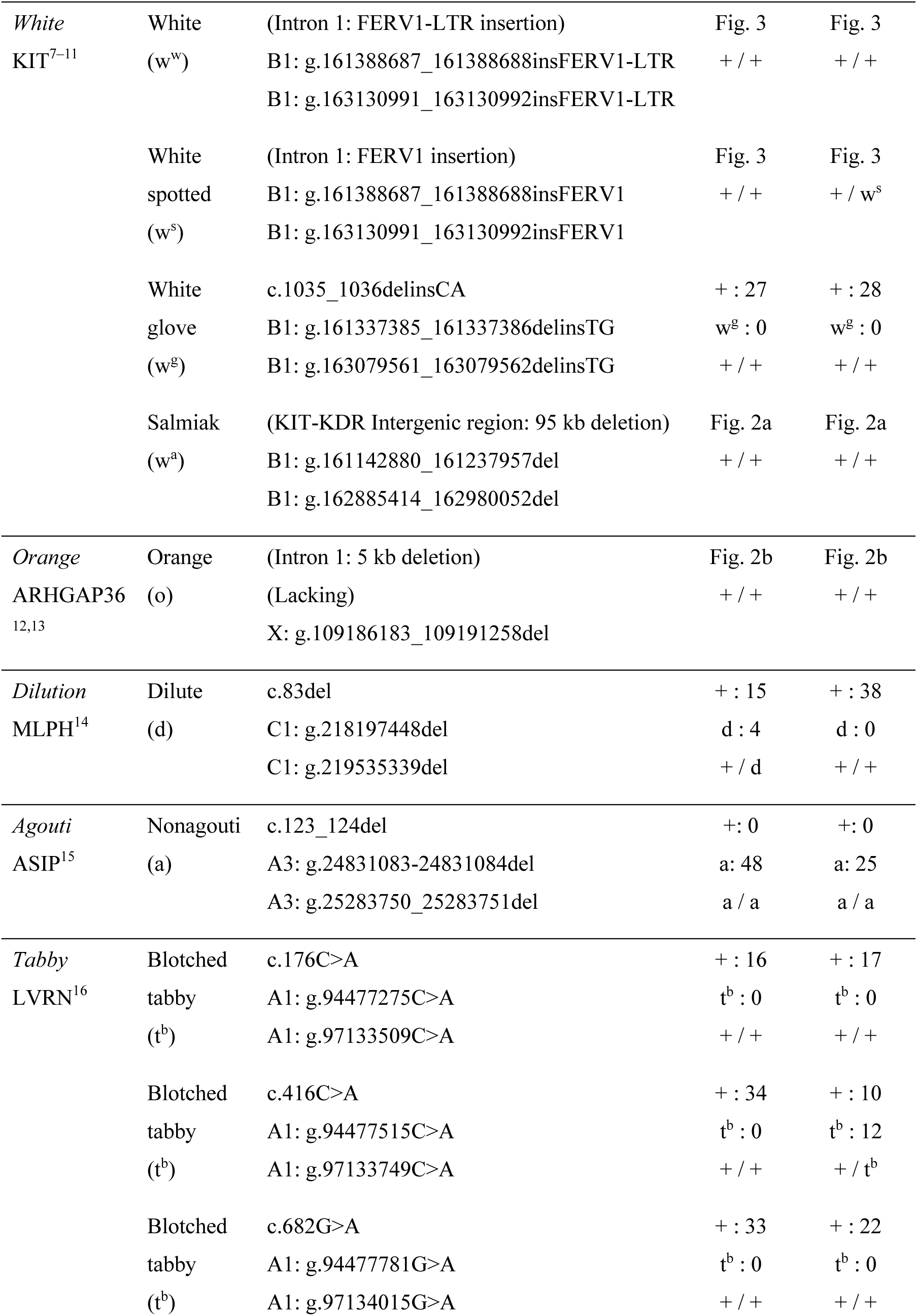
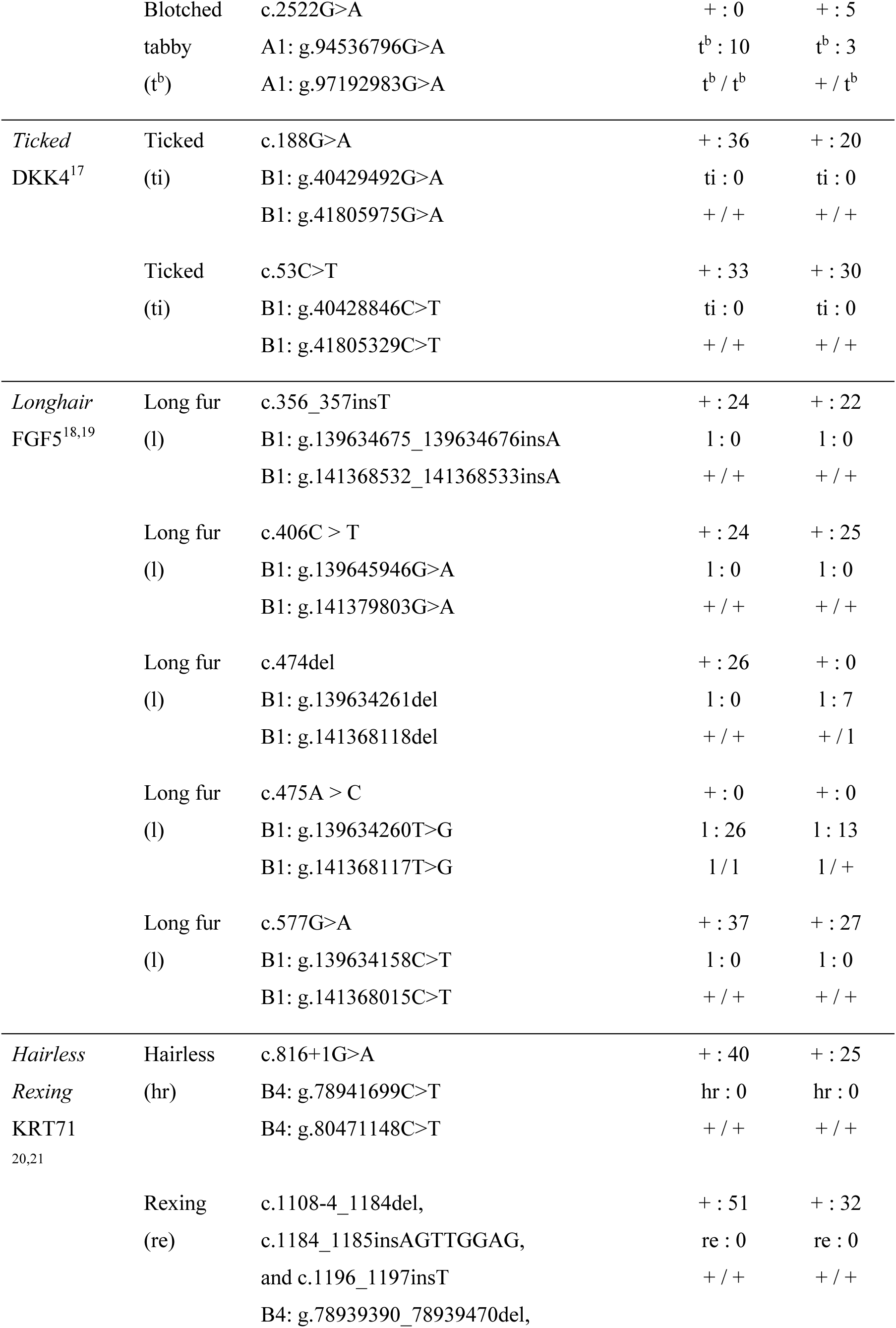
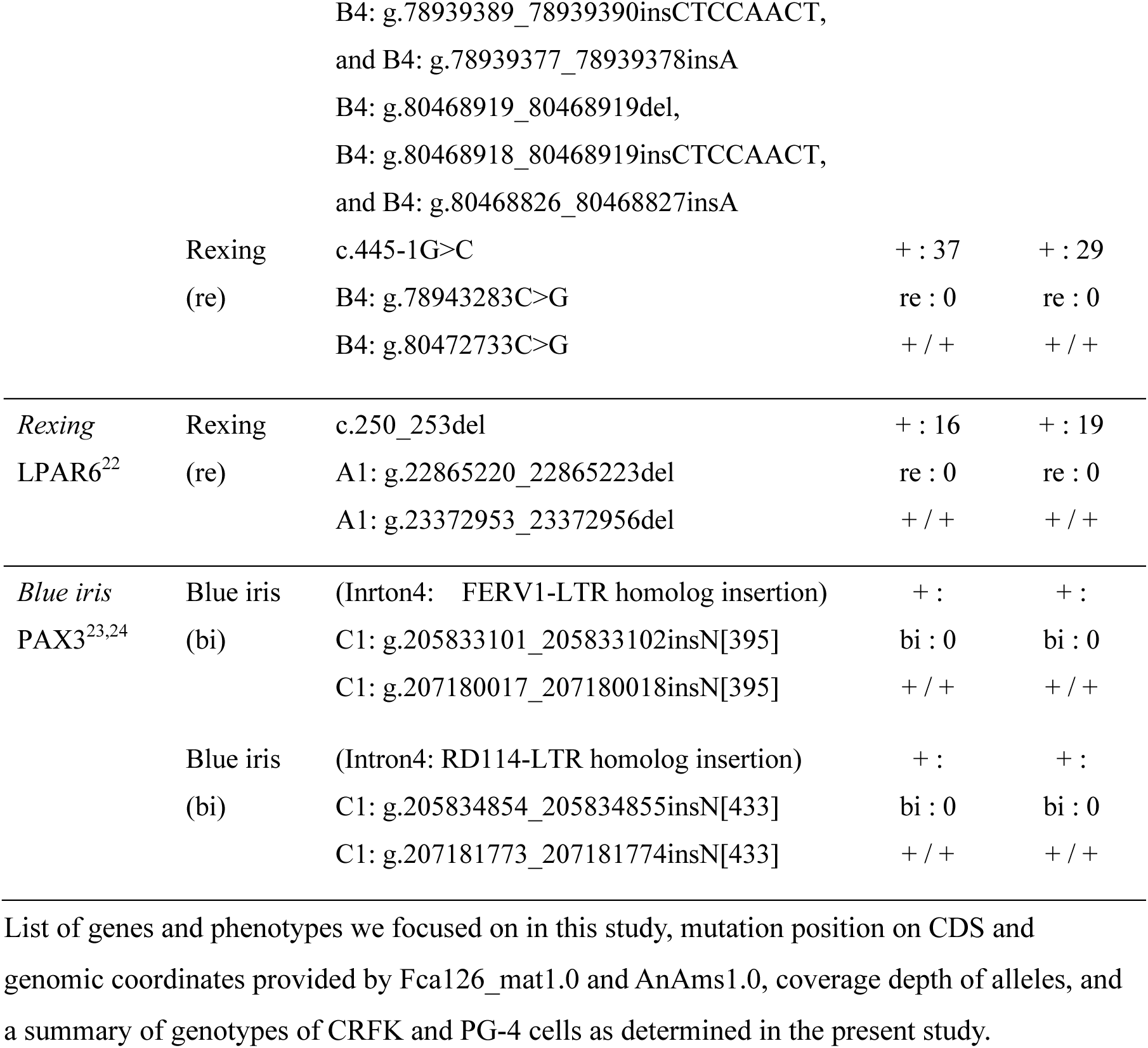
Phenotypic traits determined by DNA variants, and genotypes of CRFK and PG-4 cells.

## Methods

### Cell lines and culture conditions

CRFK cells (JCRB9035, RRID: CVCL_2426) and a feline sarcoma-positive leukemia-negative (S+L−) astrocyte cells termed PG-4 (S+L-) (JCRB9125, RRID: CVCL_3322) were obtained from the JCRB Cell Bank (National Institutes of Biomedical Innovation, Health and Nutrition, Osaka, Japan). CRFK cells were cultured in Dulbecco’s Modified Eagle Medium (Gibco, Thermo Fisher Scientific, Waltham, MA, USA) supplemented with 10% fetal bovine serum (FBS) (Gibco), penicillin (100 IU/mL), streptomycin (100 ng/mL) (Sigma-Aldrich, St. Louis, MO, USA), and 1% non-essential amino acids (FUJIFILM Wako Pure Chemical Corporation, Osaka, Japan) at 37 °C and 5% CO₂. PG-4 cells were cultured in McCoy’s 5A Modified Medium (Gibco) supplemented with 10% heat-inactivated FBS (Gibco), penicillin (100 IU/mL), and streptomycin (100 ng/mL) (Sigma-Aldrich) at 37 °C and 5% CO₂. For cell passaging, cells were washed using BASIC DPBS (no calcium or magnesium; Gibco) and then treated with TrypLE™ Express Enzyme (1X; Gibco) for 5 min at 37 °C and 5% CO₂ conditions.

### Paired-end short-read whole-genome sequencing

Genomic DNA was extracted from logarithmically growing cells (approximately 1×10⁶ cells) using the DNeasy Blood & Tissue Kit (QIAGEN, Hilden, Germany) according to the manufacturer’s protocol. The concentration and purity (A260/A280 ratio) of the extracted DNA were measured using a NanoDrop 2000c Spectrophotometer (Thermo Fisher Scientific). Genomic DNA extracted from CRFK cells and PG-4 cells was used for library preparation and paired-end sequencing (150 bp read length) at Novogene (Beijing, China) using a Rapid Plus DNA Lib Prep Kit for Illumina V2 (Illumina) and an Illumina NovaSeq X Plus platform (Illumina). Library quality control was performed by Novogene using qPCR and fragment size analysis according to Novogene’s standard protocols. The raw sequence data were obtained as .fastq files.

### Mapping of paired-end reads and quantification

Adapter sequences were trimmed, and low-quality reads were removed from the original .fastq files using fastp software (ver. 0.21.0) with default parameters. Reads in processed .fastq files were mapped to the reference genome sequence of *Felis catus*, F.catus_Fca126_mat1.0 (RefSeq assembly GCF_018350175.1) or AnAms1.0 (GenBank assembly GCA_013340865.2)^27^ using bwa mem software (ver. 0.7.19) with the setting “-R “@RG¥tID:sample1¥tSM:sample1¥tLB:lib1¥tPL:ILLUMINA”” and default parameters otherwise. Mapping results were obtained as .sam files. Text data in .sam files were transformed to binary data in sorted .bam files using Samtools (ver. 1.7) with default parameters. Index files were created for the obtained .bam files in the form of .bam.bai files.

### Small variations in coat morphology-related genes

The variations of coat morphology-related phenotypes of the donor cats from which CRFK and PG-4 cell lines originated were inferred based on the comparisons between the wild-type sequences and mapped sequences on variant positions of each coat morphology-related gene (Table 1). Coat phenotype-related polymorphisms in *TYR*, *TYRP1*, *MC1R*, *KIT*, *MLPH*, *ASIP*, *LVRN*, *DKK4*, *FGF5*, *KRT71*, and *LPAR6* genes, which relate to phenotypes such as the color, brown, red, white glove, dilution, agouti, tabby, ticked, long-fur, hairless, and rexing (Table 1), were examined using alignments of paired-end sequence reads with wild-type sequences.

### Large deletions in intron 1 of ARHGAP36 and the intergenic region between *KIT* and *KDR* genes

The deletion of a part of intron 1 of the *ARHGAP36* gene (X: 109,186,183_109,191,258 in AnAms1.0; Table 1) was reported to produce orange coat color^12,13^, and that of the intergenic region between the *KIT* and *KDR* genes (B1: 161142880_161237957 in F.catus_Fca126_mat1.0, Table 1) produces salmiak coat color^11^. The occurrence of these deletions in CRFK and PG-4 was estimated by comparing the mapped read number distributions between these regions and their neighboring regions with the same region length. It should be noted that F.catus_Fca126_mat1.0 has a deletion of the genomic region corresponding to X: 109186183_109191258 in AnAms1.0 (Table 1). Thus, AnAms1.0 (or felCat9^25^) was used as the reference genome for analysis of the orange locus.

### Insertions of FERV1-LTR and RD114-LTR into *PAX3*

The insertions of homologous sequences of the long terminal repeat (LTR) of the feline endogenous retrovirus 1 (FERV1), termed FERV1-LTR, into a specific region in intron 4 of *PAX3* (C1: g.205833101_205833102 in F.catus_Fca126_mat1.0, Table 1) or the LTR of RD114 retrovirus, termed RD114-LTR, into another specific region in the same intron (C1: g.205834854_205834855 in F.catus_Fca126_mat1.0, Table 1) are associated with blue iris color in cats.^23,24^ The occurrence of such insertions was estimated by extracting the genome reads that were mapped in both FERV1-LTR and intron 4 of *PAX3* or both RD114-LTR sequences and intron 4 of *PAX3*. The genome sequence of RD114-LTR was obtained from the NCBI database^55^.

### Long-read whole-genome sequencing

Genomic DNA of CRFK cells and PG-4 cells was extracted for long-read genome sequencing using the NucleoBond HMW DNA kit (TaKaRa, Tokyo, Japan), and libraries for the Oxford Nanopore Technologies (ONT, UK) sequencer were constructed using a ligation library preparation kit (SQK-LSK114, ONT). All procedures were performed according to the manufacturer’s instructions, and the quality and molecular weight of the genomic DNA were measured using the Qubit fluorometric quantification system (Thermo Fisher Scientific). Sequencing was conducted using a PromethION sequencer (PromethION 2 Solo, ONT) and R10.4.1 flow cells. POD5 files were basecalled to .fastq files using MinKNOW software v25.05.14.

### Insertions of FERV1 homolog into *KIT* examined by long-read sequencing

The insertion of FERV1 into a specific region of intron 1 of the *KIT* gene (B1: 161388687_161388688 in F.catus_Fca126_mat1.0, Table 1) and that of FERV1-LTR make the coat color of cats white-spotted and white throughout, respectively ^8^. The occurrence of such insertions in CRFK and PG-4 cells was estimated by mapping the reads of long-read sequencing to the genome sequence named KIT-FERV1 sequence using minimap2 (ver. 2.30), where the KIT-FERV1 sequence is a part of the *KIT* intron 1 sequence with the insertion of the FERV1 sequence, as shown in Supplementary Figure 1 of the study by David et al^8^.

### Transcriptome data analysis of coat phenotype- and iris color-related genes in various tissues of domestic cats

Mapped RNA-seq data on the F.catus_Fca126_mat1.0 were obtained in .bam and .bam.bai formats for various tissues from the Ensembl public database^56^, specifically, the file transfer protocol (ftp) site^57^. The annotation file of the F.catus_Fca126_mat1.0 in gene transfer format (GTF) was obtained from the NCBI database (Bethesda, MD, USA). In the file, the chromosome numbers were converted from NC_058368.1, NC_058369.1, etc. to A1, A2, etc.

Using FeatureCounts with the options --M-O--fraction,^58^ read counts data for each gene described in the GTF file were obtained from the .bam and .bam.bai formatted files. This count data could also be obtained from a file “F.catus_Fca126_mat1.0.ENA_gene_exp_mat_by_FeatureCounts.csv” deposited in F.catlas^59^. From these counts, the transcript level of each gene was calculated by dividing the read count by the total exon length. The total transcript level was defined as the sum of all the transcript levels of the genes. Based on this result, the transcripts per million (TPM) of each gene were calculated as 1,000,000*(transcript level)/(total transcript level).

## Results

### Short- and long-read whole-genome sequencing

For CRFK and PG-4 cells, the total output of raw sequence data of 150 bp paired-end sequencing was 60.6 Gb and 60.1 Gb, and the [average] ± [standard deviation] of mapping depth of the reads for F.catus_Fca126_mat1.0 were 23.59 ± 9.79 and 23.45 ± 8.17, respectively; the total output of raw sequence data of long-read sequencing was 9.6 Gb and 11.8 Gb, N50 values were 23.5 kb and 38.2 kb, and the [average] ± [standard deviation] of mapping depth of reads for F.catus_Fca126_mat1.0 were 3.91 ± 3.11 and 4.79 ± 3.34, respectively. The average and standard deviations specified above were estimated through the entire genome region without the regions with singular sequences where the read coverage exhibited >100.

### Predicted phenotype of the CRFK cell donor

The mapping of paired-end reads from the whole-genome sequences of the CRFK cell revealed that most coat morphology-related genes were conserved (Table 1). However, the following variations were identified in mapped genome reads from CRFK cells (Table 1): a homozygous mutation of c.123_124del of the *ASIP* gene (*nonagouti* allele), a homozygous mutation of c.475A>C of the *FGF5* gene (*long coat* allele; Fig. 1), a homozygous mutation of c.2522G>A of the *LVRN* gene (*blotched-tabby* allele), and a heterozygous mutation of c.83delT of the *MLPH* gene (*dilute* allele). No 95 kbp deletion was detected in the intergenic region between the *KIT* and *KDR* genes, nor a 5 kbp deletion was observed in intron 1 of the *ARHGAP36* gene (Fig. 2). Additionally, no genome reads were identified that could be simultaneously mapped to both the *PAX3* intron 4 and FERV1-LTR or to both the *PAX3* intron 4 gene and RD114-LTR sequences. Furthermore, four reads from the long-read sequencing mapped to the KIT-FERV1 sequence; however, these reads contained no insertions of FERV1 and FERV1-LTR (Fig. 3).

**Figure 1:**
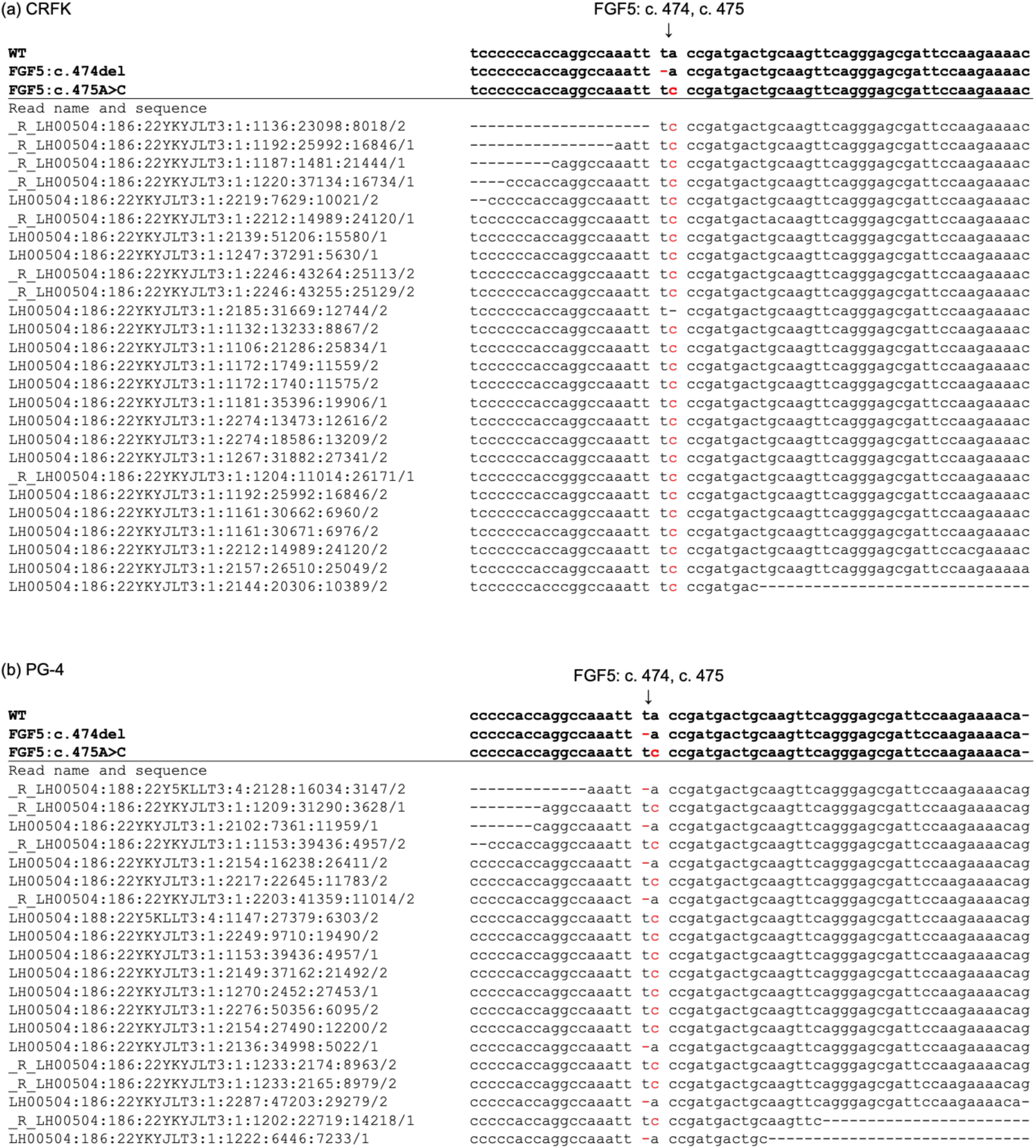
Alignment results of whole genome sequencing reads of CRFK and PG-4 cells with WT and mutant sequences of FGF5 genes. Alignment results of genome sequencing reads around FGF5: c.474 and c.475; CRFK (a) and PG-4 (b) cells. Left string indicates the name of sequenced read, and red colored symbols indicate coat phenotype-related mutations

**Figure 2:**
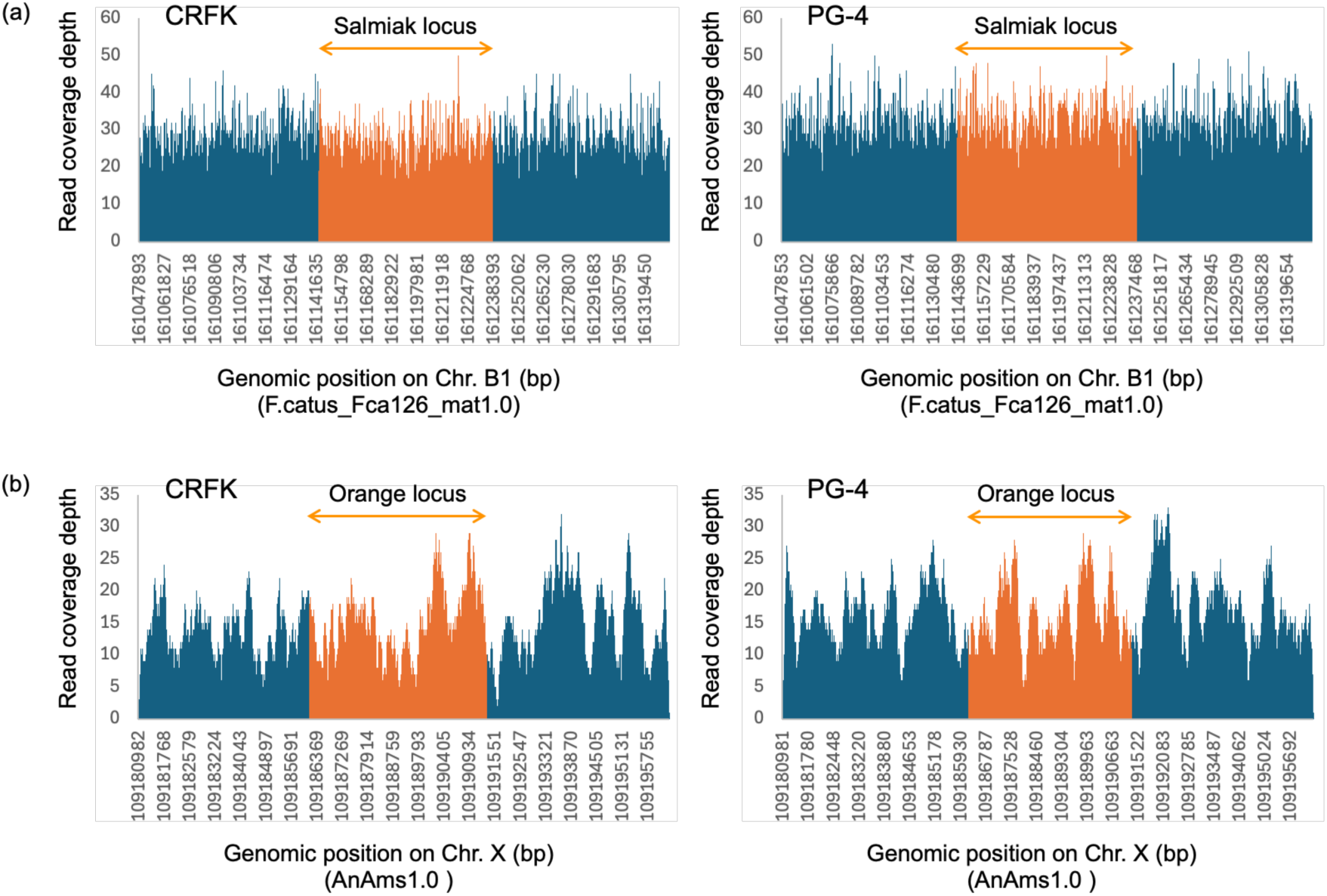
Read coverage depth distributions of CRFK and PG-4 cells around the salmiak and Orange loci. Read coverage depth distributions from 95 kbp upper stream to 95 kbp lower stream of the salmiak locus (a), and those from 5.1 kbp upper stream to 5.1 kbp lower stream of the Orange locus (b) of CRFK and PG-4 cells. Bar graph colored orange indicates coverage depth of reads mapped to the salmiak (a) and Orange loci (b), respectively. In all distributions, no notable declines in read coverage depth were observed in both salmiak and Orange loci, indicating that both cats from which the CRFK and PG-4 cells express no salmiak and orange coat.

**Figure 3:**
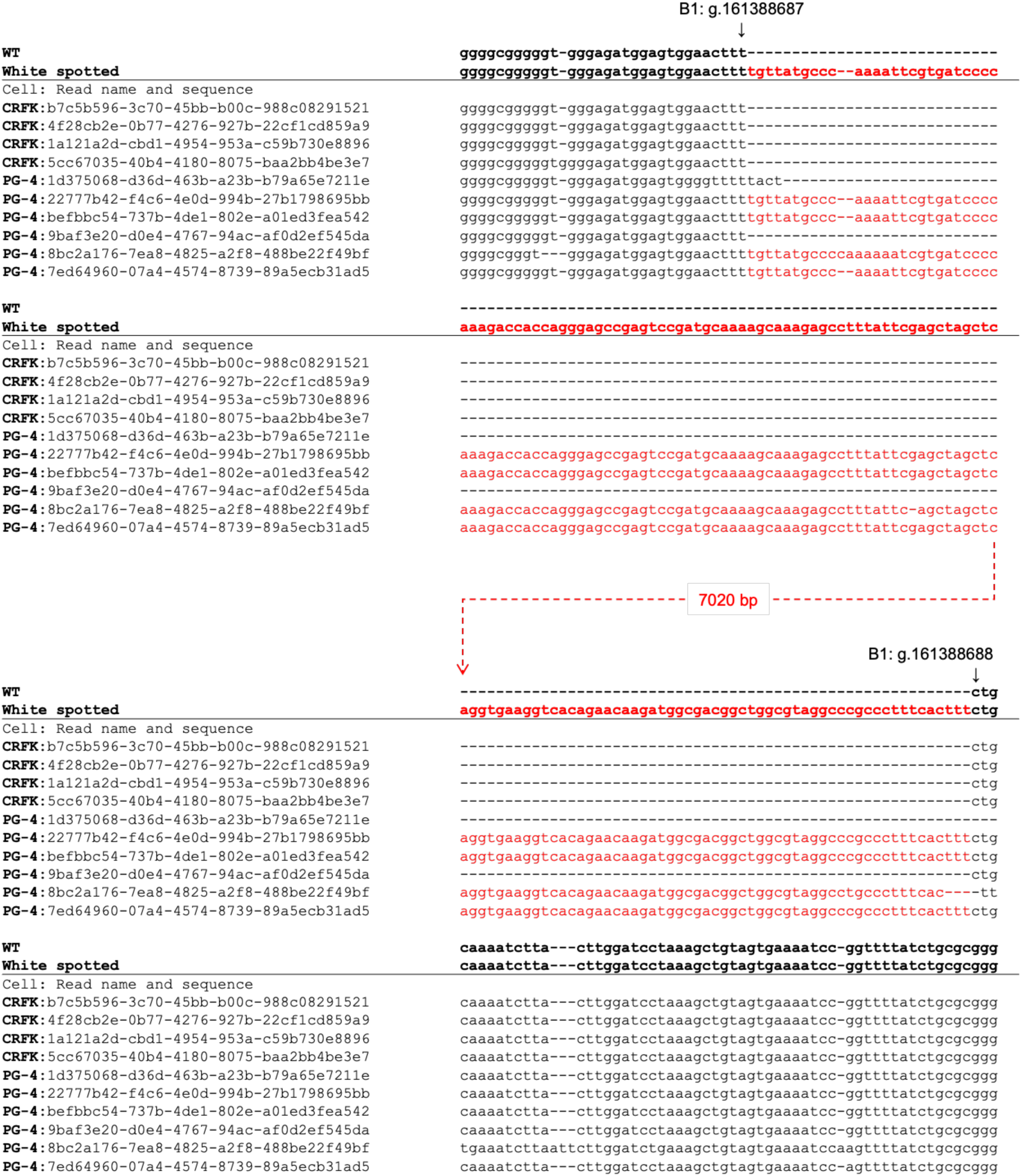
Parts of upper and lower streams of alignment results of long-read sequencing reads of CRFK and PG-4 cells with WT and KIT-FERV1 sequences of KIT intron 1. Alignment results of long-read genome sequencing reads from CRFK and PG-4 cells to the KIT-FERV1 that contains sequence around the region B1: g.161388687_161388688 (F.catus_Fca126_mat1.0). Left string indicates [the name of cell (CRFK or PG-4): name of sequenced read], and red colored sequences indicate FERV1 and aligned sequences with it. Four long reads from CRFK could align, but no reads contained the FERV1 homologue, while 6 reads from PG-4 could align, and four of the six reads contained the FERV1 homologue. This figure shows only a part of the results at the upper and lower streams of the KIT-FERV1 sequence. See Table S1 for the alignment result with the full-length KIT-FERV1 sequence

The abovementioned homozygous mutation of the *FGF5* gene was known to exhibit long fur^18^. However, the mutation of the *LVRN* did not influence coat morphology in this case because of the mutation of the *ASIP* gene that suppresses the stripe pattern on the cat’s skin coat, and the mutant genotype of the *MLPH* gene was recessive.^1^ Therefore, the cat from which the CRFK cells originated is expected to have had long, black, unstriped fur and non-blue eyes (Fig. 4).

**Figure 4:**
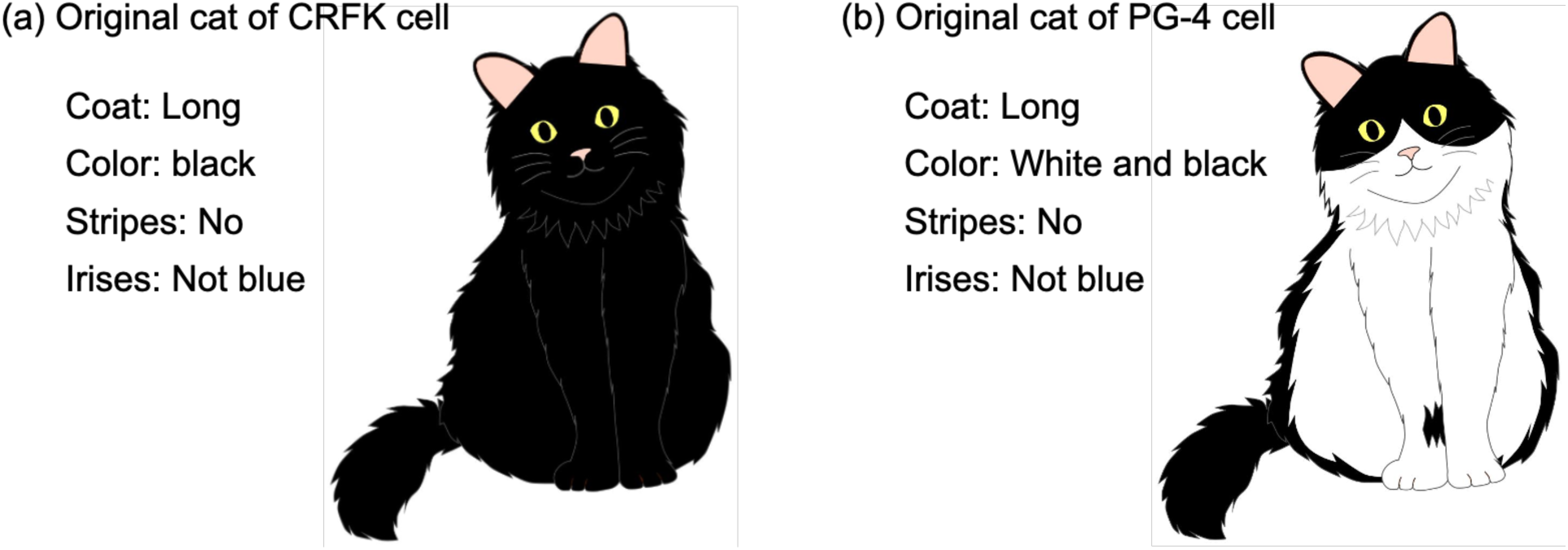
Example illustrations of inferred original cats of CRFK and PG-4 cells. An example illustration of the inferred phenotypes of coat morphologies and iris colors of the original cats of the CRFK cell (a) and the PG-4 cell (b). CRFK cells are expected to originate from a cat with long, black fur, lacking stripes, and with non-blue irises. PG-4 cells were expected to be derived from a cat with long, white and black bicolored fur without stripes and non-blue irises

### Predicted phenotype of the PG-4 cell donor

Mapped paired-end reads from the whole-genome sequences of the PG-4 cell indicated that most coat morphology-related genes were conserved but revealed the following variations (Table 1): a homozygous mutation of c.123_124del of the *ASIP* gene, a compound heterozygous mutation of c.474delT and c.475A>C of the *FGF5* gene (Fig. 1) where one allele exhibits c.474delT and the other allele exhibits c.475A>C as mentioned below, the heterozygous mutations of c.416C>A and c.2522G>A of *LVRN* gene, and a heterozygous mutation of c.904G>A of the *TYR* gene (*Siamese-color* allele). No 95 kbp deletion was identified in the intergenic region between the *KIT* and *KDR* genes, nor a 5 kbp deletion was found in intron 1 of the *ARHGAP36* gene (Fig. 2), as was similarly observed with CRFK. Additionally, we did not identify any genome reads that mapped concurrently to both the *PAX3* intron 4 and FERV1-LTR or to both the *PAX3* intron 4 gene and RD114-LTR sequences.

Notably, the genome sequence reads obtained from PG-4 cells containing bases 474 and 475 of the CDS of the FGF5 gene revealed that seven reads exhibited only a deletion at base 474, while the remaining 13 reads showed only a substitution at base 475 (Fig. 1). Accordingly, we expected a compound heterozygous mutation in the FGF5 gene of PG-4 cells. Similarly, PG-4 cells exhibited heterozygous substitutions at bases 416 and 2522 of the CDS of the LVRN gene. However, owing to the considerable distance between the genomic locations of these two bases, none of our sequencing data, including the long-read sequencing data, produced reads that contained both bases together. Therefore, we were unable to conclude whether the LVRN gene in PG-4 cells harbors a compound heterozygous mutation.

Six long-read sequences mapped to the KIT-FERV1 sequence, with four of them containing the insertions of the FERV1 homolog sequence exceeding 7 kbp (Fig. 3). This indicates that the PG-4 genome contains the heterozygous insertion of the entire FERV1 homolog sequence.

The mutant genotype of the *TYR* gene was recessive, and in this case, the two heterozygous mutations of the *LVRN* did not influence coat morphology because of the *ASIP* gene mutation, the same as in CRFK^1^. However, the effect of *FERV1* is known to be dominant^8^. Additionally, the compound heterozygous mutation of the *FGF5* gene produces long fur^18^. Therefore, the PG-4 cells likely originated from a cat with white and black (white spotting), unstriped, long fur (Fig. 4).

Notably, certain cats with white spotting may have blue eyes or atypical eye coloration; however, this occurrence is less common than that in cats exhibiting the Dominant White allele in the KIT gene^8^. Additionally, no insertions of FERV1-LTR and RD114-LTR in intron 4 of the PAX3 gene were observed. Therefore, the PG-4 cells likely originated from a cat with non-blue irises (Fig. 4).

### Expression features of coat phenotype- and iris color-related genes in various tissues

The analysis of publicly available RNA-seq data from various organs of domestic cats in the Ensembl database (https://ftp.ensembl.org/pub/data_files/felis_catus/F.catus_Fca126_mat1.0/rnaseq/) revealed that coat phenotype-related genes are expressed in various organs at levels (TPM) comparable to or greater than those in the skin (Fig. 5, Table S2). For example, *TYR*, *TYRP1*, and *MLPH* were highly expressed in the ear tip, retina, and optic nerve, and *MC1R*, *KIT*, *DKK4*, and *FGF5* were highly expressed in the brain. *KIT*, *DKK4*, and *LPAR6* tend to be expressed throughout the body, with prominent expression in the reproductive organs. Additionally, *ARHGAP36* was highly expressed in the spinal cord, and *ASIP*, along with *MLPH*, was highly expressed in the lungs. The iris color-related gene *PAX3* was also highly expressed in the cerebellum, in addition to visual organs, and, along with *DKK4*, in the kidney and embryonal tissue.

**Figure 5:**
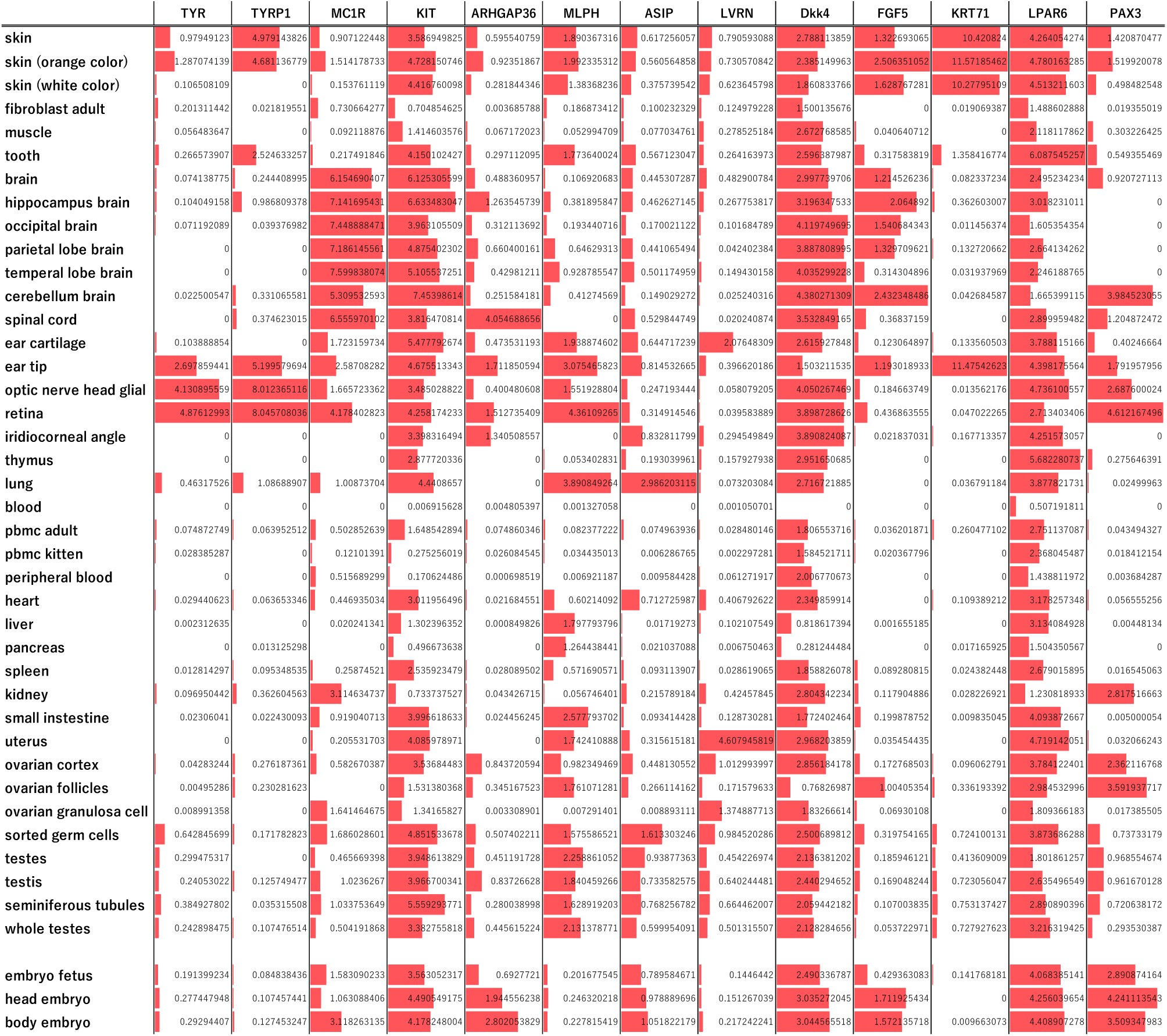
Expression levels of coat phenotype-related and iris color-related genes in various tissues. The expression level of each gene in each tissue is quantified as log_2_(transcripts per million+1). The length of the red bar for gene A for each tissue is proportional to the [expression levels of gene A in each tissue] / [the maximum expression level of gene A among all focused tissues]. Most coat phenotype-related genes were found to exhibit similar or higher expressions in various tissues other than skin, and the iris color-related gene (PAX3) was also found to exhibit similar or higher expressions in a variety of tissues other than the opt-functional organs

## Discussion

We performed whole-genome sequencing of CRFK, a cultured feline kidney cell line, and PG-4, a feline fetal astrocyte-derived cell line, and analyzed the genome data to determine the coat color, hair length, coat pattern, and iris color of the donor individuals. The results suggested that the CRFK cell line originated from a cat with long black hair without stripes and non-blue irises, and the PG-4 cell line came from a cat with long fur, black and white bicolor coat without stripes, and non-blue irises.

Coat morphology and iris color are phenotypes that can be readily identified and used to predict genotypes. However, the expressions of these genes are not limited to the skin and eyes; rather, they are expressed in various tissues throughout the body (Fig. 5). Therefore, differences in these genotypes may contribute to disease susceptibility in specific organs. The effects of these genetic variations are likely weaker than those of the rearing environment and may not be significant, especially in young, healthy individuals. However, as domestic cats have become longer-lived in recent years, individual-specific risks may become more apparent, and the underlying genotypes may exert increasing effects as the cats age. Therefore, elucidating the relationships between genotypes underlying readily recognizable phenotypes and disease risk is crucial from the perspectives of both medical care and welfare, such as prevention and symptom suppression/alleviation.

Results from research on cultured cells to elucidate disease mechanisms and develop therapeutic drugs must be interpreted with caution, i.e., not only on the experimental results obtained using established, immortalized cells but also with regard to their genotype and expression profiles of the genes harboring polymorphisms in various tissues and organs. However, so far, such information has not been available, even for certain frequently used cell lines such as CRFK. The present study and genome sequence data provide such information and facilitate a more comprehensive interpretation of research on feline cultured cell lines.

Furthermore, with the availability of detailed genomic information, genome editing can be applied to these cell lines to establish various derivative lines that differ only in specific aspects, e.g., regarding coat morphology or other traits, and compare their properties. This is expected to advance research on the regulatory mechanisms of coat color, for example, the validation of the role of the novel *KIT* isoform caused by FERV1-LTR insertion in the formation of white coat^60^, as well as the relationships between the visually identifiable phenotypes examined in the present study and disease risk and treatment methods. Simultaneously, this comprehensive genomic information can help to develop guidelines for establishing cell lines with various disease-associated genotypes using genome editing. The development of these derivative cell lines may provide valuable insights into the mechanism underlying deafness in cats with white fur and blue irises, as well as elucidate the influence of other coat-related factors in organs beyond the skin or functions unrelated to coat formation.

The limitations and future directions of this study are outlined below.

First, notably, the findings of this study are specific to the cell types used. The cells used in this study were CRFK cells (JCRB9035, RRID: CVCL_2426), which were prepared from CRFK (ATCC: CCL 94) and PG-4 (S+L-) cells (JCRB9125, RRID: CVCL_3322) obtained from PG-4 (S+L-) cells (ATCC: CRL-2032). Consequently, these cells were expected to have similar characteristics to those of the CRFK and PG-4 cells used in numerous previous studies. For example, however, the number of chromosomes has been reported to vary from 36 to 38 in CRFK^33,54^. Additionally, the karyotype of PG-4 cells, which has not yet been reported, likely differs from the normal domestic cat genome. Therefore, using the same cell line may still produce results that differ slightly from those used by other researchers. Investigating additional variation among cells in these lines remains an important future consideration.

As mentioned previously, the karyotype of CRFK differs from that of the normal domestic cat genome^33,54^. Additionally, the specific chromosomes that cause aneuploidy remain unidentified. However, identifying large structural variations using only whole-genome analysis of the present sequence read data, approximately 60 Gb paired-end sequencing data and 10 Gb long-read data, was challenging. Similarly, accurately identifying sequences and locations of various long repeat sequences, endogenous retroviral sequences, and their homologs was challenging owing to limited read coverage, particularly for long-read coverage. However, these sequences may also be associated with various diseases, as demonstrated by the involvement of FERV1-LTR in deafness in cats^8,23,24^ and the positive and negative roles of enFeLV-LTR and enFeLV-env in resistance to exogenous feline leukemia virus^61–63^. The increased volume of ongoing long-read sequencing, along with the draft genome assembly using these data, is expected to address these challenges in the future.

This study focused on coat and iris phenotypes in domestic cats. However, major histocompatibility complex (MHC) polymorphisms are strongly correlated with breed in domestic dogs^64–68^. Although research on MHC polymorphisms in domestic cats is ongoing^69–71^, the relationship with breed remains unclear. In this study, we showed the results of the primary analysis of the small indels and SNPs in MHC class I and II regions of CRFK and PG-4 cells described by Variant Call format (VCF) ^72^ (Table S3). However, because the MHC gene cluster comprises a complex and large genomic region, it is difficult to identify the donor breeds of CRFK and PG-4 using only the sequence data and analytical methods presented in this study. However, as mentioned above, further long-read sequencing and its data analysis may enable breed estimation of these cell lines.

In addition, the present genome sequence data also included mitochondrial DNA data. In this study, we presented the results of the primary analysis of the small indels and SNPs in the mitochondrial DNA of CRFK and PG-4 cells described by VCF^72^ (Table S4^72^). Although the correlations are not substantially strong, an association may exist between the polymorphisms of mitochondrial DNA and breeds of domestic cats^73–76^. Therefore, a comprehensive analysis of this data may also contribute to breed estimation.

These analyses are also critical future challenges.

## Conclusions

We performed whole-genome sequencing of CRFK and PG-4 cells and inferred the phenotype of the donors of these cells. We suggested that CRFK cells originated from a cat with long, black fur lacking stripes and non-blue irises; PG-4 cells originated from a cat with long, bicolored white and black fur without stripes, and with non-blue irises. Additionally, analysis of publicly available RNA-seq data confirmed that genes associated with coat phenotype and iris color are expressed in the skin and eyes, as well as in various other organs, indicating that variants of these genes, which affect coat phenotype and iris color, may influence physiological functions throughout the body. These insights may facilitate a more accurate interpretation of data derived from feline cultured cells and inform guidelines for developing cell lines with domestic cat genotypes that exhibit various phenotypes using genome editing techniques. This will help to elucidate the role of various coat phenotype-related and iris color-related factors in organs other than the skin and functions independent of coat formation, including the effects of melanocyte destruction caused by KIT gene mutations on the nervous system.

## Declarations

### Ethics approval and consent to participate

Not applicable.

### Consent for publication

Not applicable.

### Availability of data and materials

These raw read data of the short and long read sequencings were deposited in the DDBJ Sequence Read Archive (DRA) with BioProject accession PRJDB37530 (DRR748391-DRR748394, DRR786948, and DRR786949). All other data generated or analyzed during this study were included in this published article (and Supplementary Information files).

### Conflicting interests

The authors have no conflicting interests to declare.

### Funding

This work was supported by Japan Society for the Promotion of Science (JSPS) KAKENHI grants (award numbers 24K01783 (T. I.) and 24K09248 (A. A.)).

### Author contributions

Conceptualization: A.A., Data curation: T.K., A.K., R.H., and T.I., Formal analysis: G.T. and R.G., Funding acquisition: A.A., Investigation: G.T., R.G., T.K., R.H., and A.A., Methodology: G.T., R.G., T.K., T.I., and A.A., Project administration: N.S. and A.A., Supervision: N.S. and A.A., Validation: R.G. and A.A., Visualization: T.G., N.S., and A.A., Writing - original draft: A.A., Writing - review & editing: T.G., T.K., T.I., and N.S.

## Acknowledgements

Not applicable.

## Figure captions

**Table S1: Alignment results of long-read sequencing reads of CRFK and PG-4 cells**

(File: Table_S1)

Alignment results of long-read sequencing reads of CRFK and PG-4 cells with WT and KIT-FERV1 sequences of KIT intron 1.

**Table S2: Gene expression levels in various tissues**

(File: Table_S2)

Gene expression levels (transcripts per million) in various tissues estimated from domestic cats (Felis catus) in the Ensembl database (https://ftp.ensembl.org/pub/data_files/felis_catus/F.catus_Fca126_mat1.0/rnaseq/)

**Table S3: Lists of polymorphisms in MHC class I and II regions of CRFK and PG-4 cells.**

(File: Table_S3.zip)

Variant Call Format (VCF)^72^ files, text files providing lists of SNPs and small indels, for MHC class I region in CRFK (a) and PG-4 (b) cells, and those for MHC class II regions in CRFK (c) and PG-4 (d) cells. These VCF files were obtained as follows: 1) Similar to “Mapping of paired-end reads and quantification” in the Methods section, .bam and .bam.bai files of the mapping results of the original paired-end short read data of the whole genome sequencing to the reference genomes were obtained. Here, genome sequencing data were mapped to the reference genomes “Felis catus FLA distal class I region genomic sequence” (GenBank: EU153402.1) and “Felis catus FLA extended class II, class II, class III, proximal and central class I region genomic sequence” (GenBank: EU153401.1) for MHC class I and II, respectively. 2) Using the “MarkDuplicates” program of Picard (ver. 2.27.4) to obtain .bam files with the identification of duplicate reads. 3) By applying the HaplotypeCaller program of GATK (ver. 4.3.0.0) to previously obtained .bam file and .fasta file of reference genomes with default parameter values, VCF files of SNPs and small indels lists were obtained.

**Table S4: Lists of polymorphisms in mitochondrial DNA of CRFK and PG-4 cells.**

(File: Table_S4.zip)

VCF^72^ files for the mitochondrial DNA in CRFK (a) and PG-4 (b) cells. VCF files were obtained similarly to those presented in Table S3 using “Felis catus mitochondrion, complete genome” (GenBank: NC_001700.1) as the reference genome.

## Notes

### Competing Interest Statement

The authors have declared no competing interest.

### Summary of Updates

Figures 2 and 4 were revised; Table 1 was revised; Some sentences in the text were revised; Supplemental files were updated.

